# Evolution of ancient satellite DNAs in extant alligators and caimans (Crocodylia, Reptilia)

**DOI:** 10.1101/2023.04.18.537305

**Authors:** Vanessa C. Sales-Oliveira, Rodrigo Zeni dos Santos, Caio Augusto Gomes Goes, Rodrigo Milan Calegari, Manuel A. Garrido-Ramos, Marie Altmanová, Tariq Ezaz, Thomas Liehr, Fabio Porto-Foresti, Ricardo Utsunomia, Marcelo de Bello Cioffi

## Abstract

Crocodilians are one of the oldest extant vertebrate lineages, which exhibits a combination of evolutionary success and morphological resilience that have persisted throughout the history of life on Earth. Such an ability to endure over such a long geological time span is of great evolutionary importance. Here, we performed a comprehensive analysis of the satellite DNA diversity of the extant alligators and caimans, making significant progress in our understanding of the evolution of repetitive regions present in ancient genomes. The alligators and caimans displayed a small number of satDNA families (varying between 3 and 13 satDNAs, in *A. sinensis* and *C. latirostris*, respectively) as well as little variation both within and between species, highlighting an exceptional long-term conservation of satDNA elements throughout evolution. We also tracked the origin of the ancestral forms of all satDNAs belonging to the common ancestor of Caimaninae and Alligatoridae. Fluorescence in situ experiments showed distinct hybridization patterns for the identical ortholog satDNAs, indicating their inner dynamic evolution. Why, in addition to their previously known low genetic, karyotype, and morphological diversity, have crocodilians altered so little over such a long period of time with such a highly variable genome fraction? We argued that such an “evolutionary package” was likely the outcome of severe demographic declines or founder events and that it represents evolutionary responses to a long-lasting bottleneck history.

## Introduction

Eukaryotic genomes are highly variable in structure and size because of the presence of vast quantities of repetitive DNA (Biémont and Vieira 2006; López-Flores and Garrido-Ramos 2012). Satellite DNAs (satDNAs) are a common component, accounting for an important part of the genome in most animal and plant genomes (reviewed by Plohl et al. 2014; Garrido-Ramos 2017, 2021). In general, a genome has a varied number of satDNA families (the satellitome, Ruiz-Ruano et al. 2016) with varying nucleotide sequences and genomic abundance (Csink and Henikoff 1998; Kuhn et al. 2008; Kuhn et al. 2010; Plohl et al. 2012; Feliciello et al. 2015; Prakhongcheep et al. 2017; Palacios-Gimenez et al. 2017). Although some examples of small arrays scattered throughout euchromatin have been documented (Ruiz-Ruano et al. 2016; Feliciello et al. 2011, 2015; Kuhn et al. 2012; Brajković et al. 2012; Larracuente 2014; Pavlek et al. 2015; Pita et al. 2017; de Lima et al. 2017; Robledillo et al. 2020; Milani et al. 2021), these sequences are often found in centromeres and in pericentromeric and subtelomeric heterochromatic areas (reviewed in Plohl et al. 2012; Garrido-Ramos 2017; Šatović-Vukšić and Plohl 2023). More than just “junk DNA” (as for a long time considered) several studies have revealed that satDNAs have a role in a variety of biological processes, including gene regulation (Joshi and Meller 2017), centromere function (Rošić et al. 2014), chromatin modulation (Ugarkovic 2005), and spatial chromosomal structure (Pathak et al. 2013; Jagannathan et al. 2018, 2019).

These sequences represent one of the fastest evolving genomic components, which often leads to high levels of interspecific sequence diversity even within closely related species, exhibiting very different profiles (both quantitative and qualitative) of satDNAs in their genomes (Henikoff et al. 2001; Plohl et al. 2012; Garrido-Ramos 2017). However, some satDNAs can persist over long periods, spanning dozens (or even hundreds) of million years (de la Herrán et al. 2001; Mravinac et al. 2002; Robles et al. 2004; Plohl et al. 2010; Lorite et al. 2017; Halbach et al. 2020; dos Santos et al. 2021). Although the direct causes for this conservation are not always clear, the acquisition of molecular function (Garrido-Ramos et al. 1995; Mravinac et al. 2002, 2005; Schueler et al. 2010; Fachinetti et al. 2015), slow rates of genomic evolution (de la Herrán et al. 2001; Robles et al. 2004) or just chance events (Camacho et al. 2022) have been proposed. The combination of cytogenetics and genomics studies has proven to be particularly useful in elucidating numerous aspects of genome evolution and organization (Graphodatsky 2007; Deakin et al. 2019), with particular emphasis on repetitive DNAs (Rovatsos et al. 2015; Ruiz-Ruano et al. 2016; dos Santos et al. 2021; Kretschmer et al. 2022, among others). Particularly, due to their tandemly repeated genomic organization, satDNA studies in non-model organisms were boosted in the late few years, especially with the development of several assembly-free pipelines designed for using raw reads (Novák et al. 2013, 2017; Harris et al. 2019; Vondrak et al. 2020). In this context, several satDNA catalogs were characterized from a variety of invertebrate and vertebrate species (Ruiz-Ruano et al. 2016; Silva et al. 2017; Kirov et al. 2018; Mora et al. 2020; de Lima and Ruiz-Ruano 2022; Peona et al. 2022; Goes et al. 2022; Kretschmer et al. 2022).

Crocodilians are one of the oldest extant vertebrate lineages, demonstrating the evolutionary success and morphological resilience that span the history of life on Earth (Brochu 2003). Extant crocodilian species have preserved physical and ecological traits for nearly 100 million years, unlike other vertebrates that have undergone significant diversity (Grigg et al. 2001; Brochu et al. 2004; Bronzati et al. 2015). Crocodilians have a key position in vertebrate phylogeny because, combined with dinosaurs, pterosaurs, and modern birds, they compose the archosaurs, a monophyletic group (Janke and Arnason 1997; Iwabe et al. 2005; Green et al. 2014). Crocodylia is classified into three families: Crocodylidae, Gavialidae, and Alligatoridae, with approximately 27 species (Uetz 2023). Alligatoridae comprises eight species divided into four genera (*Alligator*, *Melanosuchus, Paleosuchus*, and *Caiman*). Except for the *Alligator* genus, where *A. mississippiensis* and *A. sinensis* are limited to the Southeastern United States and China, respectively, all the other six species are presently found in South America, being more widespread in Brazil (Brochu 2003; Oaks 2011).

The karyotypes of all current Alligatoridae species were recently revised using conventional differential staining and up-to-date molecular cytogenetic approaches (Oliveira et al. 2021). Although there is a limited amount of diversity and certain level of karyotype stasis (with diploid numbers equal to 2n=42 and 2n=32 for all Caimaninae (*Caiman, Paleosuchus,* and *Melanosuchus*) and Alligatorinae (*Alligator)* species, respectively), their genomic content revealed significant interspecific divergence.

Here, we performed a comprehensive analysis of the diversity of satellite DNAs of Alligatoridae. We isolated, characterized, and compared the full catalogs of satDNA families (i.e., the satellitomes) of 5 out of 8 extant Alligatoridae species by integrating genomic and chromosomal data. The results revealed strong sequence conservatism among Caimaninae species with very limited diversity of their satDNA library. Furthermore, fluorescence in situ assays in all extant Alligatoridae species showed that the identical satDNA orthologs can exhibit various hybridization patterns, indicating their high evolutionary dynamics.

## Results

### Bioinformatic satDNA characterization

After several iterations (*C*. *yacare* = 5, *C*. *latirostris* = 7, *M*. *niger* = 4, *P*. *trigonatus* = 4, and *A*. *sinensis* = 2), we characterized 39 satDNAs in Alligatoridae, where repeat unit lengths ranged from 23 (PtrSat11-23) to 6317nt (ClaSat02-6317) and the average of their A+T content was 46.9%. Specific features of the satDNAs in each species are summarized in **Table 2.** The number of iterations performed for each species was a consequence of the results obtained in each round so that when no new tandem repeats were discovered in a given round the analysis was not continued. Thus, for example, in the case of *A. sinensis* no tandem sequences were discovered in the third interaction. Iterations using RepeatExplorer2 after TAREAN did not return any characterized satellite DNA for the five species analyzed.

In general, alligators and caimans analyzed here exhibited few satDNA families (average=7.8 ± 4.08 satellite per species) and also a small diversity in the within-and between-species level, intraspecific cases with similarity greater than 50% and less than 80% were classified as the same superfamily, while interspecific cases with similarity greater than 50% were placed in the same Group, divided here into four different Groups. Based on sequence alignments, three main groups of satDNAs were identified showing at least 50% of similarity that encompassed satDNAs from at least four species, named here as Group 1 (N=19 satDNAs shared among Caimaninae), Group 2 (N=8 satDNAs shared among Caimaninae) and Group 3 (N=6 satDNAs shared among Alligatoridae) (**Figure 1**). In addition, a fourth group included sequences from two species (ClaSat04-536 and PtrSat09-490), while the remaining 4 satDNAs did not show any similarity with other sequences (**Table 2**). This classification helped us to delimit the origin of some satDNAs and follow their diversification patterns in each species.

**Figure 1.**
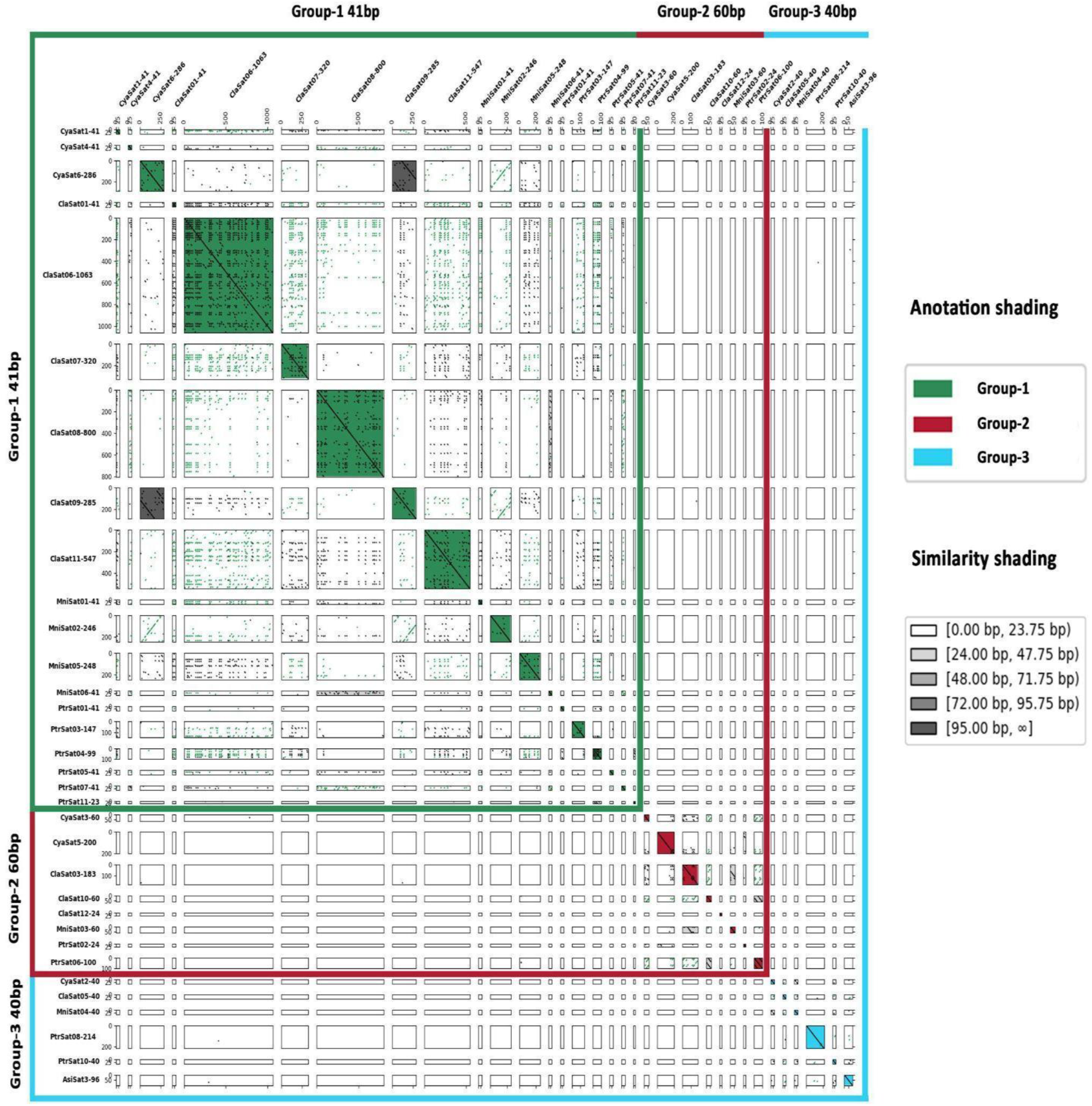
All-against-all dotplot of Caimaninae satellite DNA, with the division of these sequences among the three groups presented in the analysis in green (group 1), red (group 2), and blue (group 3), and the similarity among each satDNA family, represented by a white to the black color ladder.

Expanding our analyses, as we found a significant RUL variability in each of those groups, we generated a global dot-plot with sequences from the above-mentioned groups (**Figure 2**). As expected, sequences belonging to the same group showed similarities as revealed by the dotplots, which also indicated that satDNAs exhibiting variable RULs within groups probably emerged from the diversification of existing repeats as HORs.

**Figure 2:**
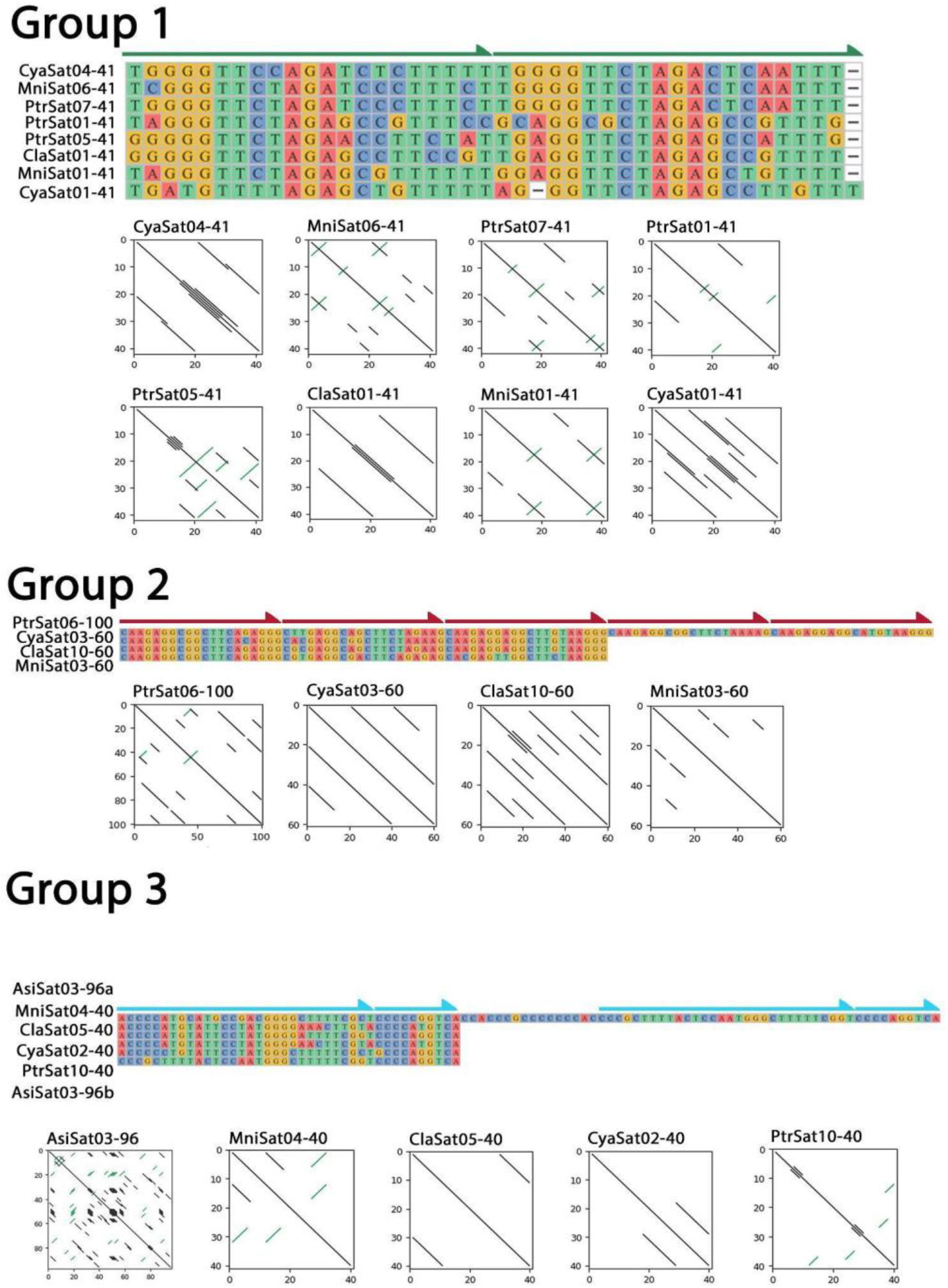
Presence of high-order repeats in the 3 groups, demarcated by arrows in alignments, representing the subunits that derived the sequences in each group. In addition, dotplots of each satDNA family, presenting the internal repetitions among all monomers, are also indicated.

We generated dotplots for each satDNA monomer, and we found that several families can be classified as higher-order repeats (HORs), indicating that the origin of novel satDNA families is usually related to the diversification of existing families in these species (**Figures 1 and 2**). For instance, in Group 1, only satDNAs with 41bp-long monomers were characterized. By analyzing them, we found that these are HORs containing two subunits (21 bp + 20 bp, **Figure 2**). Considering Group 2, monomers with 60 bp are observed in *Caiman* and *Melanosuchus*, while in *P. trigonatus*, monomers exhibit 100 bp. Our analysis revealed that 60 bp-long monomers are HORs containing 3 subunits (20bp + 20bp + 20bp, **Figure 2**). Finally, Group 3 is the only example of shared satDNAs between all the analyzed species here. Repeat monomers of 40 bp are predominant among these satDNAs, and dot-plot analysis revealed a heterogeneous HOR composed of two different subunits (29 bp + 11 bp, **Figure 2**). Remarkably, in *A. sinensis*, a 96 bp-long satDNA was characterized in this group and its monomer sequence is composed of two 40-bp units interconnected by an unrelated 16-bp sequence, thus being a complex HOR composed of two heterogeneous HORs and an intervening sequence (40bp + 16bp + 40bp, **Figure 2**).

BLAST searches against the genome of *A. sinensis* revealed that satDNAs classified as groups 1 and 2 were not found in this species, while significant matches were observed for sequences belonging to group 3 (CyaSat02-40, ClaSat05-40, MniSat04-40, PtrSta10-40 and AsiSat03-96) and group 4 (ClaSat04-536 and PtrSat09-490). Also, significant matches were produced against ClaSat02-6317 and ClaSat13-398 (results are summarized in Supplemental Table S2). These results indicate that groups 1 and 2 of sequences emerged after the split of Caimaninae and Alligatorinae, while groups 3 and 4 are shared among the representatives of both subfamilies. In addition, Alligatorinae-specific AsiSat01-1717 and AsiSat02-60 satDNAs returned abundant significant matches, as expected.

ClaSat02-6317 and ClaSat13-398 were highly abundant (**Supplemental Table S2**) and their origin could be traced, but each one exhibited different patterns in BLAST. While ClaSat13-398 produced multiple hits against *A. sinensis* genome (n=1712), the largest alignment corresponded to the size of the monomer (398 bp), indicating that this sequence is potentially shared among all Alligatoridae species. On the other hand, ClaSat02-6317 produced even higher hits (n=6264), but the largest observed alignment was 1073 bp, indicating a significant change in monomer size along the history. Remarkably, the obtained TSI for ClaSat13-398 was low (TSI=0.21), suggesting that this sequence is dispersed along the genome of *A. sinensis*, which is also observed for the Group 4 satDNA ClaSat04-536 (TSI=0.25). While ClaSat02-6317 exhibited a higher TSI in *A. sinensis* (TSI=0.74), we hypothesize that this is due to its larger monomer size. Since the fragments of paired-end sequencing are usually around 300-400 bp and the monomer of this satDNA is >6,000bp, the obtained TSI is most likely due to mapping in the same monomer, not mapping in adjacent monomers.

Collectively, our analyses revealed, for *A. sinensis*, that: i) satellite DNAs classified as group 4 show a dispersed pattern throughout the *A. sinensis* genome, as evidenced by TSI and FISH; and ii) hybridization signals of AsiSat03-96 (group 3) were not visible, probably due to the organization in short arrays (exhibited high TSI, but low number of alignments in BLAST). This is in contrast to the alligator-specific satellites that appear clustered at loci on long arrays, consistent with results obtained in FISH experiments, high TSI and a large number of alignments in BLAST.

### Chromosomal location of satDNAs with differential abundance between species

We analyzed the chromosomal location of SatDNAs that were successfully amplified by PCR belonging to group 1 (ClaSat01-41; ClaSat06-1063; ClaSat07-320; ClaSat08-800 and ClaSat11-547), group 2 (ClaSat10-60), group 3 (ClaSat05-40), group 4 (ClaSat04-536) in addition to the two exclusively ones found in *A. sinensis* (AsiSat01-1717 and AsiSat02-60) in all Alligatoridae species to check their chromosomal distribution. Additionally, the ungrouped satellites ClaSat02-6317 and ClaSat13-398 were tested but none of them gave a FISH signal in any species (data not shown).

Concerning the ClatSatDNAs, except for the satellite ClaSat04-536 (group 4), which showed no hybridization signals in any species, all other SatDNA sequences were found in (peri-) centromeric heterochromatin regions in all Caimaninae species **(Figures 3-5)**. Both alligators (*A. sinensis* and *A. mississippiensis*) showed no hybridization signal for any of the ClaSatDNAs investigated (data not shown). Here to illustrate, we present the results for representative selected ClaSatDNAs from each of the major groups identified **(Figures 3–5)**. The satDNA ClaSat01-41, belonging to Group 1 (the most frequent group present in each species) were mapped in two chromosomal pairs in all Caimaninae species except *P. trigonatus,* which did not display any hybridization signal **(Figure 3)**. However, despite sharing the same motifs, some divergent and species-specific chromosomal location patterns were observed among ClaSatDNAs from Group 1 among species **(Supplemental Fig. S1–S2)**. For satellites in groups 02 and 03, numerous chromosomal pairs containing these sequences were found in nearly all species **(Figures 4 and 5)**. *M. niger* is distinctive for displaying hybridization signals on only two chromosomal pairs for group 3 satellites **(Figure 5d)**.

**Figure 3.**
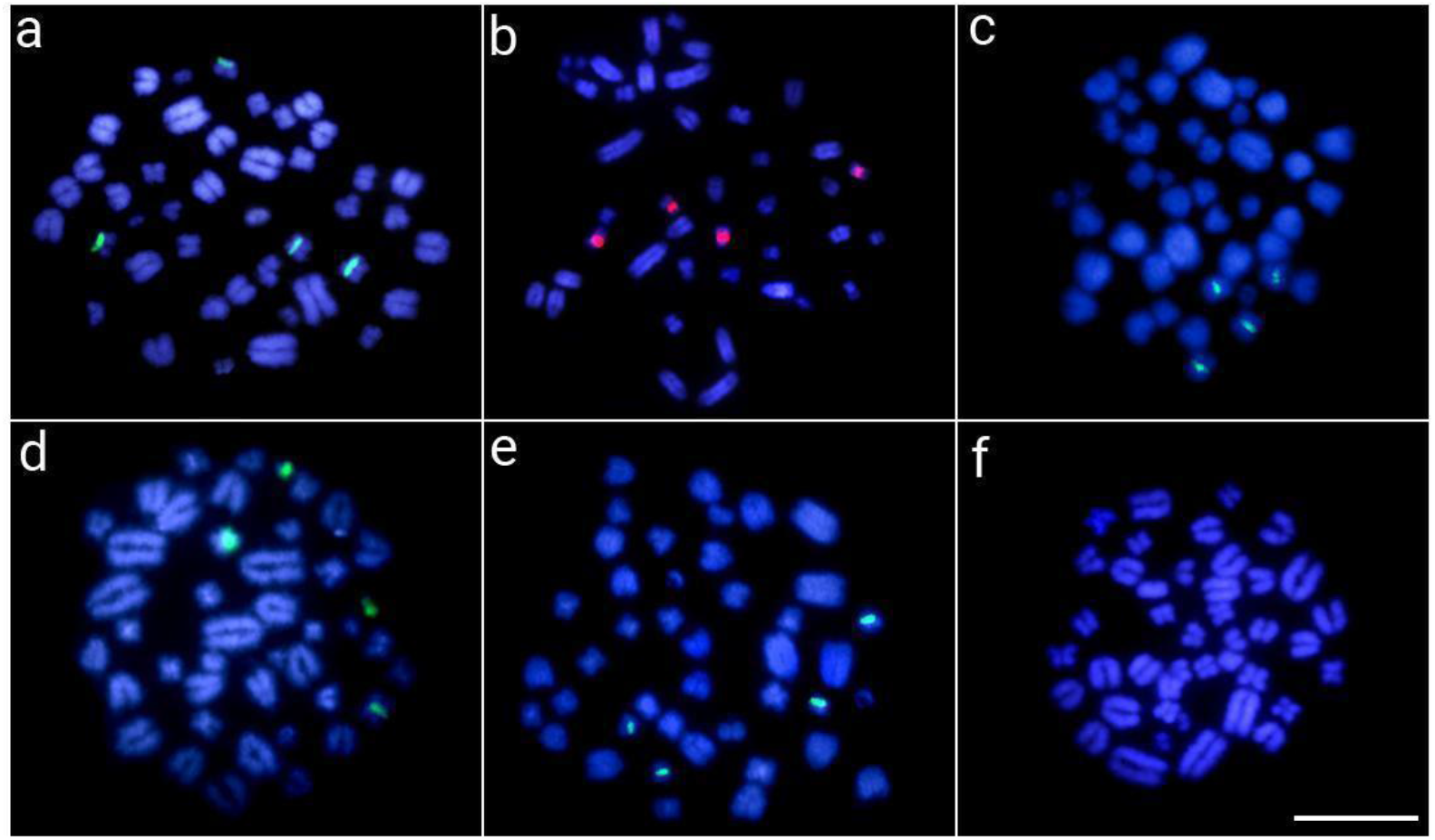
Metaphase chromosomes from *C. crocodilus* (a), *C. latirostris* (b), *C. yacare* (c), *M. niger* (d), *P. palpebrosus* (e) and *P. trigonatus* (f) after in situ mapping of ClaSat01-41 (group 1). Chromosomes were counterstained with DAPI (blue). Scalebar=20μm

**Figure 4.**
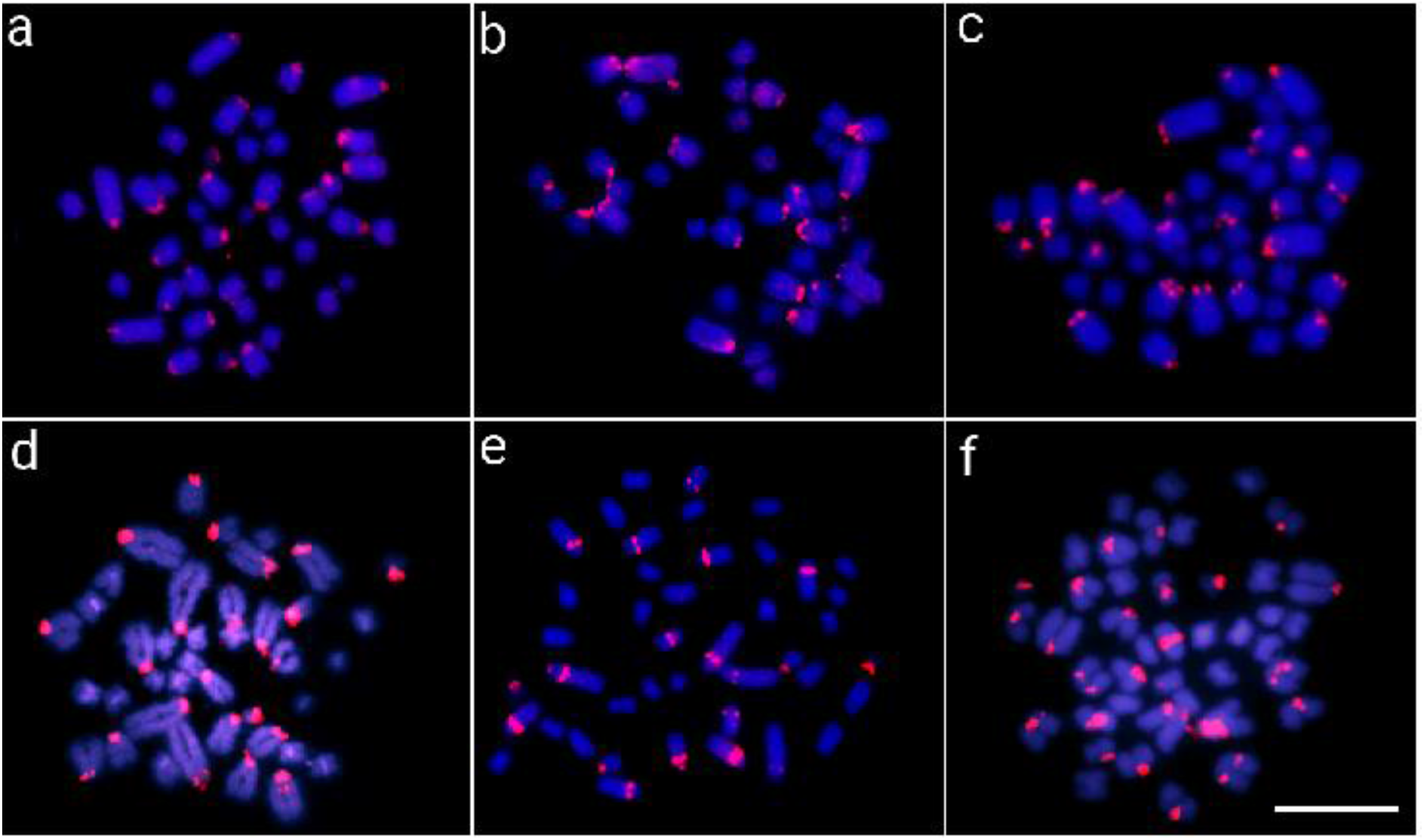
Metaphase chromosomes from *C. crocodilus* (a), *C. latirostris* (b), *C. yacare* (c), *M. niger* (d), *P. palpebrosus* (e) and *P. trigonatus* (f) after in situ mapping of ClaSat10-60 (group 2). Chromosomes were counterstained with DAPI (blue). Scalebar=20μm

**Figure 5.**
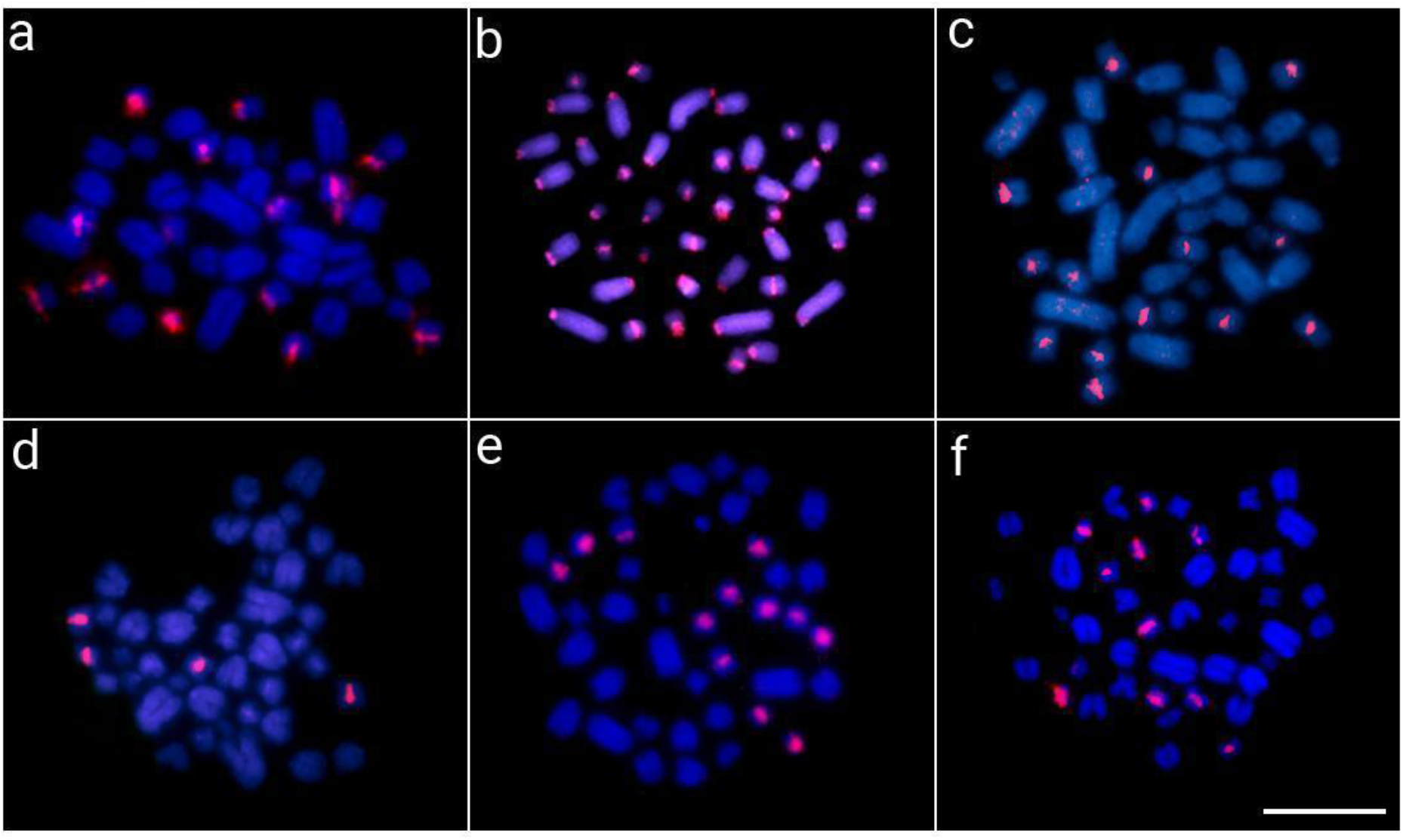
Metaphase chromosomes from *C. crocodilus* (a), *C. latirostris* (b), *C. yacare* (c), *M. niger* (d), *P. palpebrosus* (e) and *P. trigonatus* (f) after in situ mapping of ClaSat05-40 (group 3). Chromosomes were counterstained with DAPI (blue). Scalebar=20μm

Besides, we also mapped the two exclusive satDNAs presented in *A. sinensis* genome (AsiSat01-1717 and AsiSat02-60) in all Alligatoridae species. Both AsiSatDNAs showed hybridization signals only in *Alligator* species. While AsiSat01-1717 was exclusively mapped in several chromosomes, AsiSat02-60 was mapped in all centromeres of both species **(Figure 6)**.

**Figure 6.**
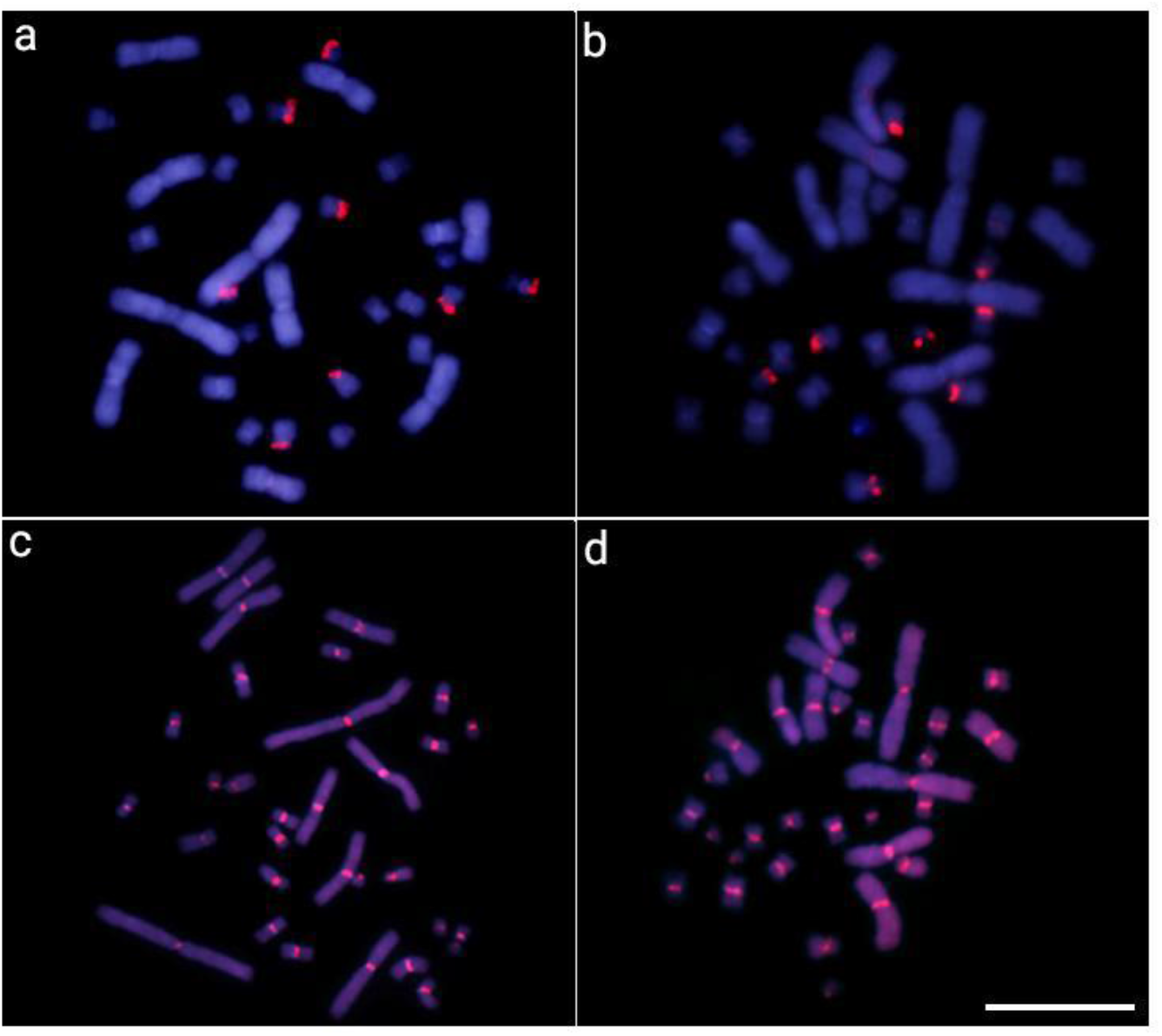
Metaphase chromosomes from *A. sinensis* (a and c) and *mississippiensis* (b and d) after in situ mapping with AsiSat01-1717 (a and b) and AsiSat02-60 (c and d) probes. Chromosomes were counterstained with DAPI (blue). Scalebar=20μm

## Discussion

Despite the fact that both alligators’ complete genomes were characterized some years ago (Wan et al. 2013; Green et al. 2014; Rice et al. 2017), genome-wide investigations of satDNAs in this group were never undertaken. SatDNAs are well known to be underrepresented in genome assemblies (Peona et al. 2021), particularly those genomes assembled using short-read sequencing technology, as is the case with alligators. In this context, knowledge about satDNAs in crocodilians was limited to individual sequences isolated by restriction endonucleases (Kawagoshi et al. 2008). Given that high-throughput satellitome analysis has been very enlightening for understanding the satDNA evolution in various organisms, we used a chromosome-and genomic-based approach to try to describe the satellitome from members of all current Alligatoridae genera for the first time. In a time span of around ∼70 Myr (million years), many satDNA sequences are shared among the species, assisting in the hypothesis that they are derived from small sequences, as shown in **Figure 2.** Furthermore, in following fluorescence in situ tests the distinct hybridization patterns for the identical ortholog satDNAs were observed.

After mining satellite DNAs using well-established bioinformatic pipelines (Ruiz-Ruano et al. 2016; Novák et al. 2017), we found that alligators’ satellitomes are among the smallest catalogs described until now, varying between 3 and 13 satDNAs, in *A. sinensis* and *C. latirostris*, respectively. Using TAREAN, two sequences were characterized in the first iteration of *A. sinensis*, one in the second, and no tandem sequences were discovered in the third interaction. We also used RepeatExplorer2 in an attempt to characterize some sequences not presented by TAREAN. Nevertheless, no novel sequence was offered, resulting in the species under study having the lowest inventory of satDNA. In recent years, several satellitomes from a wide range of species, including plants and animals, have been identified (Ruiz-Ruano et al. 2016, 2019; Camacho et al. 2022; Goes et al. 2022; de Lima and Ruiz-Ruano 2022; Montiel et al. 2022; Peona et al. 2022). These investigations showed that satellite DNA profiles are very dynamic. For example, characiform fish satellitomes display a significant quantitative and qualitative variation, with some species exhibiting a few dozen (Goes et al. 2022), while others can show more than one hundred satDNAs (Utsunomia et al. 2019). Here, we found that all alligators are similarly satDNA-poor constituting a common trend in this group.

Novel satDNA families can emerge by variable mechanisms and from multiple genomic regions, like introns, transposable elements, and/or existing satDNA families (Ruiz-Ruano et al. 2018; Valeri et al. 2018; Vondrak et al. 2020). Our data showed a generally low intraspecific satDNA diversity, suggesting that most new satellite sequences arose from previously existing ones. For instance, *C. latirostris* exhibited 13 satDNAs, but six and three of them were grouped in groups 1 and 2, respectively (**Table 2**). Interestingly, this low diversity is also reflected on the interspecific level, and more than 90% of the 39 satDNA families described for Alligatoridae can be grouped into 4 groups of sequences. After their origin, multiple subsequent events of diversification, like higher-order repeats (HORs) formation, shaped the evolution of satellite DNAs in Alligators. While groups 1 and 2 were characterized as regular HORs, group 3 sequences are characterized as heterogeneous ones, as their sequences were derived from a repetition of two subunits of 29 bp and 11 bp **(Figure 2).** In addition, longer satellites of each group have also undergone a complex evolutionary process in which different cycles of amplification and divergence have followed one another, including even the inclusion of intervening sequences between established HORs. It is very common to find that new longer satDNAs have arisen from the complex diversification of shorter satDNAs families already existing in the genome (de la Herrán et al. 2001; Navajas-Pérez et al. 2005; Ruiz-Ruano et al. 2019).

The long-term evolution of satellite DNA catalogs in related species can be explained by the library hypothesis. Fundamentally, it states that changes in the profiles of sat DNAs among species are mostly quantitative in the “library”, rather than multiple *de novo* origins (Fry and Salser 1977). Here, we could track the origin of the ancestral forms of satDNAs belonging to groups 1–4 to, at least, the common ancestor of Caimaninae (Groups 1, 2, and 4) and Alligatoridae (Group 3). We found a significant similarity of satDNAs among species, while only four were species-specific. The long-term maintenance of satDNAs is notable. In this context, the conservation could be related to the acquisition of cellular function (Fry and Salser 1977), particular genomic organization (dos Santos et al., 2021), or slow rates of evolution (de la Herrán et al. 2001). Previous studies found slow rates of molecular evolution within crocodilians (Green et al. 2014); thus, we hypothesize that satDNAs also evolved slowly in this group (as discussed below). In squamate reptiles, satDNAs generally have a homogeneous composition and a prevalent centromeric chromosomal location (Capriglione 2000). While the great majority of sequences are of recent origin and only observed in closely-related related species (Capriglione et al. 1994; Rudykh et al. 1999; Ciobanu et al. 2003, 2004; Grechko et al. 2005), several (and most common ones) are largely conserved in unrelated species (Olmo et al. 2002).

The chromosomal mapping analysis revealed that all characterized satellites showed the general same chromosomal location (i.e., large peri-and centromeric blocks) among species, showing specific patterns for each one **(Figures 3–6 and Supplemental Fig. S1–S2)**. On the other hand, it is interesting to see that group 1 satellites, even being the most abundant in the Caimaninae genome, show a visible block of FISH signal in only two chromosomal pairs. When using the FISH technique, as a specific satDNA sequence can actually display a variety of array structures (dispersed and/or clustered into long and nonrandom arrangements) among species, it results in a range of labeling patterns at the chromosomal level. This is particularly true, for example, for the Group 3 ClatSat05-40 because, although being abundant in the genome of *A. sinensis,* it exhibits non-cluster organization (as indicated by our BLAST results in **Supplemental Table S2**), which hindered in situ experiments from producing any detectable hybridization signals at chromosomal level. We hypothesize that this could well explain the FISH patterns observed in Caimaninae for group 1 satDNAs, although we cannot verify this as we do not have the complete sequence of their genomes nor are these satDNAs present in the

### A. sinensis genome for comparisons

Although the presence of multiple dispersed loci composed of a single copy or a few tandem copies of a satDNA family is a fact today (Šatović-Vukšić and Plohl 2023), the accumulation of satDNAs (as well as other repetitive DNA families) in centromeres and in heterochromatic regions is characteristic, as observed in many other groups (Stornioli et al. 2021; Iwata et al. 2013; Shang et al. 2010; Garrido-Ramos 2017; Šatović-Vukšić and Plohl 2023). In Alligatoridae, constitutive heterochromatin is restricted to the pericentromeric regions (Oliveira et al. 2021) and includes the (peri)centromeric clusters found for the satDNAs isolated in this study. It can be assumed that, as in other species, some of these satellites would be part of the centromeric chromatin. Such colocalization (i.e., the tendency to occupy similar locations on non-homologous chromosomes) might have been facilitated by the reunion of centromeres at the first meiotic prophase bouquet (Mravinac and Plohl 2010; Ruiz-Ruano et al. 2016). This is especially true in Caimaninae since the karyotypes of all species are dominated by acrocentric chromosomes. Besides, considering their ubiquitous presence in the (peri)centromeric region of all analyzed Caimaninae species, it is supposed that satDNA must play at least some important role in the centromeric organization. There are many examples of species in which different satDNAs occupy different (peri)centromeric regions (reviewed in Garrido-Ramos 2017, 2021). In this context, the existence of large and small chromosomes in Caimaninae could be favoring the structural differences at the (peri)centromeric level between different chromosomes (Lanfredi et al. 2001). Both *Paleosuchus* species stood out by presenting differentiated hybridization patterns from the rest of the Caimaninae. In this sense, we have previously observed in comparative genomic hybridization (CGH) and rDNA mapping analyses that both *Paleosuchus* species highlighted a small sharing of repetitive sequences with another cofamiliar species, demonstrating a distinctive pattern (Oliveira et al. 2021).

Both alligators (*A. sinensis* and *A. mississippiensis*) displayed hybridization signals only for two (AsiSat01-1717 and AsiSat02-60) of all the satDNAs investigated **(Figure 6)**. Furthermore, AsiSat02-60 was exclusively mapped in all centromeres of both *Alligator* species. That is, these two species have conserved the same (peri)centromeric satDNA in all their chromosomes highlighting its possible important role in the centromeric and pericentromeric organization, a role that it may be sharing with AsiSat01-1717 in some chromosomes. Alligatorinae long diverged (∼70 Myr) from all the other Caimaninae and, have highly rearranged karyotypes (2n=32) that are predominantly metacentric, in contrast with all Caimaninae species that have 2n=42 chromosomes and karyotypes dominated by acrocentric chromosomes (reviewed in Oliveira et al. 2021). We have proposed that 2n=32 represents the likely ancestral state and that the karyotype diversification in Caimaninae was followed by a series of Robertsonian rearrangements in which centric fissions played a key role (Oliveira et al. 2021). Accordingly, alligators’ satellitomes are among the smallest catalogs described until now for any species, with only 3 satDNAs identified.

The potential significance of satDNA sequences in centromeric function has been extensively explored. In this instance, one would expect that the same satDNA would be conserved across all examined species. While the same satellite has been conserved in centromeres of Alligatorinae species for about 70 Myr, the chromosomal rearrangements that have taken place in the Caimaninae lineage would have caused the emergence and diversification of new satellite DNAs that have replaced them in the (peri)centromeric regions. Some of them, such as those of group 3, were already present in a dispersed form in the ancestral genome of Alligatoridae, as was possibly the case with the satDNAs of group 4 and the ungrouped satDNAs ClaSat02-6317 and ClaSat13-398 (**Supplemental Table S2**), still dispersed in all Alligatoridae species. In fact, everything points to epigenetic control of the centromeric function and to a structural role of satDNAs indifferent to its sequence (reviewed in Garrido-Ramos 2015, 2017, 2021) but in which its capacity to acquire non-B DNA conformations would be fundamental (Kasinathan and Henikoff 2018). Thus, the maintenance and/or replacement of some satDNAs by others would be a stochastic process (Camacho et al. 2022). In the case of Alligatoridae, possibly affected in turn by the slow evolution of their genomes. Extant crocodiles have limited rates of morphological (Mook 1927; Sill 1968), molecular (Green et al., 2014), and karyotype diversification (Cohen and Clark 1967; Cohen and Gans 1970; Oliveira et al. 2021). Here, we added a new piece to this puzzle by demonstrating that the present-day satellitome (particularly the Caimaninae species) shares common satDNA libraries among its species, despite their long time of divergence. But, why have they also changed so little in such a highly variable genome fraction over such an enormous span of time? Would such low genetic, karyotype, and morphological variability be related to the low number of extant crocodilian species?

Crocodylomorpha (a category that comprises living and extinct crocodilians) first appeared roughly 250 million years ago, and its 27 existing species are among the biggest living ectothermic animals. As a result, their survival over such a long geological time span is of great evolutionary importance. They do, however, have a rich fossil history that includes hundreds of extinct species, revealing a hidden past of incredible variety and complexity (Mannion et al. 2019; Stubbs et al. 2021). Oaks (2011) has questioned the traditional notion of crocodiles as old “living fossils,” arguing that most extant crocodilians are remnants of formerly successful lineages in terms of diversity and range. Crocodylomorpha is the only pseudosuchians to have survived the Triassic-Jurassic (TJ) extinction event, which happened around 200 million years ago (Brusatte et al. 2008; Toljagic and Butler 2013). Furthermore, after the mid-Miocene climatic optimum, there was a huge drop in crocodilian diversity, which coincided with global cooling and glacial advancement. During the Pliocene, the number of taxa is believed to have decreased from around 26 to 8, representing the greatest extinction rate over the previous 100 million years (Markwick 1998). As a result, the selection of an “evolutionary package” with similar genomic, chromosomal, morphology, and physiology to what is currently observed among extant species most likely resulted from drastic demographic declines or founder events and represented evolutionary responses to a long-term bottleneck history.

## Methods

### Samples, DNA extraction, and chromosomal preparation

**Table 1** summarizes the collecting sites, number, and sex of individuals used in this investigation. The sampling is similar to that previously examined by Oliveira et al. (2021). In vitro blood cultures were used to obtain chromosomal preparations (Viana et al. 2016, Johnson Pokorná et al. 2016). The usual phenol-chloroform-isoamyl alcohol procedure was used to extract genomic DNA (gDNA) from blood stored in 100% ethanol (Sambrook and Russel 2001). Blood samples were collected from free-living South American animals with the permission of the environmental agencies ICMBIO/SISBIO (License no 71857-7) and SISGEN (ABFF266). *Alligator sinensis* blood samples were obtained from lawfully housed animals in Europe (CITES certificate numbers EU 0228-1057/14, ES-CC-00041/07C, ES-CC-00036/07C, 50721-18, DE-DA190814-5, DE-DA190814-6). There were no major injuries to the animals, and all free-living individuals were returned to their respective collecting sites.

**Table 1.** Species, sample size (N), sex, and locality of the analyzed individuals.

| Species | N | Locality/Origin of Samples |  |
| --- | --- | --- | --- |
| *Caiman crocodilus*<br>(Spectacled caiman) | 2♀, 2♂ | Amazonas (BR)<br>(Amazon Basin) | 3°22'34.7" S<br>60°19'20.7" W |
| *Caiman latirostris*<br>(Broad-snouted caiman) | 4♀, 6♂ | São Paulo (BR)<br>(Cerrado) | 22°33'53.1" S<br>48°00'35.2" W |
| *Caiman yacare*<br>(Yacare caiman) | 2♀, 8♂ | Mato Grosso (BR)<br>(Pantanal) | 16°19'32.0" S<br>57°46'35.7" W |
| *Melanosuchus niger*<br>(Black caiman) | 2♀, 2♂ | Amazonas (BR)<br>(Amazon Basin) | 3°25'50.4" S<br>66°02'35.0" W |
| *Paleosuchus palpebrosus*<br>(Cuvier's dwarf caiman) | 3♀, 3♂ | Pará (BR)<br>(Amazon Basin) | 1°18'19.7" S<br>48°19'05.0" W |
| *Paleosuchus trigonatus*<br>(Schneider's smooth-fronted caiman) | 3♀, 4♂ | Amazonas (BR)<br>(Amazon Basin) | 3°06'52.0" S<br>60°01'58.0" W |
| *Alligator mississippiensis*<br>(American alligator) | 2♀, 2♂ | Canberra University collection<br>(Australia) |  |
| *Alligator sinensis*<br>(Chinese alligator) | 4♀, 2♂ | Private collections<br>(Germany) |  |

### Sequencing Data

Two broad-snouted caimans *C. latirostris* and the Schneider’s smooth-fronted caiman *P. trigonatus* were selected for low-pass shotgun sequencing on the BGISEQ-500 platform at BGI (BGI Shenzhen Corporation, Shenzhen, China), yielding 2.76 Gb, 2.76 Gb and 2.67 Gb, respectively. Raw reads are available in the Sequence Read Archive from the NCBI (SRA-NCBI) under the accession numbers: SRR19901397 *(C. latirostris* male), and SRR19901398 (*C. latirostris* female), SRR19901554 (*P. trigonatus* female). To search and compare satDNAs in other Alligatoridae species, we also collected genomic data available in the SRA-NCBI for the Yacare caiman *Caiman yacare* (SRR1609243), the black caiman *Melanosuchus niger* (SRR1609245) and for the Chinese alligator *Alligator sinensis* (SRR953089), thus encompassing all the extant Alligatoridae genera. The general features of sequencing data are summarized in Supplemental Table S1.

### Satellite DNA characterization and comparative analyses

After gathering sequencing data for all the species as mentioned earlier, we performed a quality (Q>30) and adapter trimming with Trimmomatic (Bolger et al. 2014) for each library separately. After that, we proceeded to the characterization of satDNAs in each species. We performed several iterations of TAREAN (Novák et al. 2017) and filtered the identified satDNAs with DeconSeq (Schmieder and Edwards 2011) following the protocol of Ruiz-Ruano et al. (2016). We analyzed 2× 500,000 reads in each iteration until no low-or high-confidence satellite DNA was found. Then, we performed additional iterations with the RepeatExplorer2 software (Novák et al. 2020), in the search for ring graphs, characteristics of the satellite DNA sequences in this analysis, due to their tandem characteristic, in an attempt to characterize some satDNA not presented in the analysis by TAREAN. After multiple iterations, we filtered and removed multigene families (5S rDNA and/or U snDNA) from the catalog. Then, we performed a similarity search among the remaining sequences with RepeatMasker using a custom python script (https://github.com/fjruizruano/ngs-protocols/blob/master/rm_homology.py), grouping them as the same sequence variant (≥ 95% of similarity), same family (≥ 80% of similarity) or different SatDNA families sharing a same superfamily (≥ 50% of similarity) in each species (Ruiz-Ruano et al. 2016). After that, we estimated the Kimura’s divergence, using Kimura 2-parameter model from the script calcDivergenceFromAlign.pl of RepeatMasker suite and abundance values for all satDNAs families with the “cross_match” option in RepeatMasker software (Smit et al. 2020), using 2× 5,000,000 reads for each library, except for *Melanosuchus niger* and *Caiman yacare*, because their libraries had fewer reads, performing the analysis with 2× 1,213,376 and 2× 1,608,245, respectively (Table 2; Supplemental Fig. S1). Genomic abundance of each satDNA was given as the number of mapped reads in each satDNA divided by the number of analyzed nucleotides. Finally, we classified each satellite family based on decreasing abundance order, as Ruiz-Ruano et al. (2016) suggested. The specific features of each satDNA family are observed in Table 2. Each catalog of satDNAs was deposited on the GenBank with accession numbers OP169024–OP169026 (*A. sinensis*), OP169027–OP169032 (*C. yacare*), OP169033– OP169038 (*M. niger*), OP169039–OP169049 (*P. trigonatus*), and OP169050–OP169062 (*C. latirostris*).

**Table 2:** General features of Alligatoridae satellitomes characterized with TAREAN. SF=superfamily, RUL=repeat unit length. Divergence per family was expressed as the percentage of Kimura divergence. SatDNAs that have 50% or more identity belong to the same Group.

| SF | satDNA | RUL | A+T (%) | Abundance (%) | Divergence (%) | TSI | Interspecific Divergence (%) | Group |
| --- | --- | --- | --- | --- | --- | --- | --- | --- |
| 1 | CyaSat01-41 | 41 | 63.4 | 2.210933 | 8.79 | 0.91 | 7.75 | 1 |
|  | CyaSat02-40 | 40 | 50.0 | 0.211760 | 8.35 | 0.98 | 8.60 | 3 |
| 2 | CyaSat03-60 | 60 | 41.7 | 0.099923 | 18.50 | 0.98 | 16.12 | 2 |
| 1 | CyaSat04-41 | 41 | 58.5 | 0.084393 | 21.64 | 0.89 | 20.40 | 1 |
| 2 | CyaSat05-200 | 200 | 43.5 | 0.061851 | 8.67 | 0.96 | 8.73 | 2 |
| 1 | CyaSat06-286 | 286 | 57.3 | 0.060993 | 4.29 | 0.61 | 2.74 | 1 |
| 1 | ClaSat01-41 | 41 | 46.3 | 1.0775 | 11.70 | 0.93 | 11.38 | 1 |
|  | ClaSat02-6317 | 6317 | 51.0 | 0.733486 | 7.94 | 0.87 | 8.34 |  |
| 2 | ClaSat03-183 | 183 | 40.4 | 0.268017 | 9.30 | 0.87 | 10.13 | 2 |
|  | ClaSat04-536 | 536 | 26.1 | 0.262798 | 4.83 | 0.92 | 4.21 | 4 |
|  | ClaSat05-40 | 40 | 47.5 | 0.207570 | 9.51 | 0.87 | 10.55 | 3 |
| 1 | ClaSat06-1063 | 1063 | 53.3 | 0.181244 | 9.09 | 0.74 | 12.78 | 1 |
| 1 | ClaSat07-320 | 320 | 53.1 | 0.129428 | 8.40 | 0.89 | 12.09 | 1 |
| 1 | ClaSat08-800 | 800 | 55.9 | 0.128862 | 14.84 | 0.99 | 6.46 | 1 |
| 1 | ClaSat09-285 | 285 | 46.5 | 0.073626 | 3.90 | 0.95 | 7.02 | 1 |
| 2 | ClaSat10-60 | 60 | 40 | 0.069188 | 17.79 | 0.4 | 14.05 | 2 |
| 1 | ClaSat11-547 | 547 | 51.7 | 0.068965 | 9.81 | 0.69 | 14.37 | 1 |
| 2 | ClaSat12-24 | 24 | 33.3 | 0.052656 | 16.21 | 0.82 | 12.10 | 2 |
|  | ClaSat13-398 | 398 | 32.7 | 0.042435 | 8.61 | 0.41 | 9.70 |  |
| 1 | MniSat01-41 | 41 | 58.5 | 1.581945 | 8.99 | 0.87 | 6.14 | 1 |
| 1 | MniSat02-246 | 246 | 55.7 | 0.442201 | 5.54 | 0.74 | 5.72 | 1 |
|  | MniSat03-60 | 60 | 40.0 | 0.240271 | 17.03 | 0.96 | 19.59 | 2 |
|  | MniSat04-40 | 40 | 52.5 | 0.110039 | 11.23 | 0.97 | 7.49 | 3 |
| 1 | MniSat05-248 | 248 | 55.6 | 0.044746 | 10.56 | 0.57 | 11.51 | 1 |
| 1 | MniSat06-41 | 41 | 56.1 | 0.032021 | 18.32 | 0.90 | 15.36 | 1 |
| 1 | PtrSat01-41 | 41 | 41.5 | 1.092207 | 05.02 | 0.91 | 4.95 | 1 |
| 2 | PtrSat02-24 | 24 | 37.5 | 0.705184 | 10.34 | 0.98 | 10.53 | 2 |
| 1 | PtrSat03-147 | 147 | 55.1 | 0.594545 | 10.02 | 0.95 | 8.95 | 1 |
| 1 | PtrSat04-99 | 99 | 51.5 | 0.332255 | 04.07 | 0.72 | 3.79 | 1 |
| 1 | PtrSat05-41 | 41 | 53.7 | 0.328567 | 12.86 | 0.92 | 9.45 | 1 |
| 2 | PtrSat06-100 | 100 | 45 | 0.251841 | 9.69 | 0.97 | 8.08 | 2 |
| 1 | PtrSat07-41 | 41 | 56.1 | 0.155107 | 16.09 | 0.96 | 16.09 | 1 |
| 3 | PtrSat08-214 | 214 | 28 | 0.139850 | 01.05 | 0.57 | 1.79 | 3 |
|  | PtrSat09-490 | 490 | 31.6 | 0.112221 | 4.29 | 0.96 | 5.24 | 4 |
| 3 | PtrSat10-40 | 40 | 45 | 0.111815 | 12.41 | 0.38 | 10.12 | 3 |
| 1 | PtrSat11-23 | 23 | 52.2 | 0.028522 | 13.70 | 0.05 | 9.42 | 1 |
|  | AsiSat01-1717 | 1717 | 50.6 | 0.234325 | 13.39 | 0.95 | 14.51 |  |
|  | AsiSat02-60 | 60 | 38.3 | 0.153713 | 3.51 | 0.99 | 3.65 |  |
|  | AsiSat03-96 | 96 | 34.4 | 0.020063 | 9.25 | 0.93 | 18.97 | 3 |

To compare the satellitomes of multiple species, we performed a similarity search with RepeatMasker (https://github.com/fjruizruano/ngs-protocols/blob/master/rm_homology.py) considering all the *de novo*-characterized satDNA sequences. Then, we aligned the monomers of all satDNAs showing at least 50% similarity with MUSCLE (Edgar 2004). In addition, we generated individual self-dotplots of the satDNA sequences and a general one with Flexidot (Seibt et al. 2018).

Taking advantage of the fact that the genomes of both species of alligators are sequenced we also conducted a BLAST search (blastn, word size=11, e-value=1e-6), comparing the entire list of satDNAs to the genome of *Alligator sinensis* (GCA 000455745.1), constructed using Illumina Hiseq2000 (Wan et al. 2013). We did not perform any structural or quantitative analysis on array sizes and/or organization because only short reads were employed for this assembly. As a result, BLAST searches provided more useful information on the presence or absence of satDNAs in the genome of *A. sinensis*. To get an estimation of the degree of tandem structure for the satDNAs in each species, we calculated the Tandem Structure Index (TSI), as demonstrated in Camacho et al. (2022). This value is calculated as the quotient of the number of paired reads mapped against a satDNA family and the total number of reads (https://github.com/fjruizruano/SatIntExt).

### Primer design and polymerase chain reaction (PCR)

We designed primer pairs for 12 satellite DNA families characterized from *C. latirostris* and two satellite DNA families characterized for *A. sinensis,* creating convergent primers for satellites larger than 1000bp and divergent primers for satellites smaller than 1000bp. We verified if those primer pairs anchors in conserved regions of monomers and used them to PCR-amplify in all Alligatoridae species. The PCRs contained 1× PCR buffer, 1.5 mM of MgCl2, 200 µM of each dNTP, 0.5 µL of each primer, 10–100 ng/µL of gDNA, and 0.2 µl of Taq DNA polymerase in a total volume of 25 µL. The PCR program included an initial denaturation at 95 °C for 7 min, followed by 34 cycles at 95 °C for 45s, 61 °C for 1 min, 72 °C for 1 min, and a final extension at 72 °C for 7 min. The PCR products were checked in 2% agarose gel.

### Probe labeling and fluorescence in situ hybridization (FISH)

Ten out of 14 satDNAs were successfully amplified, and the PCR products were labeled with Atto550-dUTP (red) or Atto488-dUTP (green) according to the manufacturer’s recommendations using the Nick-Translation mix kit (Jena Bioscience, Jena, Germany). The probes were then hybridized in all other Alligatoridae species according to the methodology reported by Yano et al. (2017). To corroborate the FISH results, at least 30 metaphase spreads were examined in each individual. Photos were obtained with CoolSNAP on an Olympus BX50 microscope (Olympus Corporation, Ishikawa, Japan), and the images were processed using Image-Pro Plus 4.1 software (Media Cybernetics, Silver Spring, MD, USA).

## Data Access

The genome sequencing data generated in this study have been submitted to the NCBI BioPro569 ject database (https://www.ncbi.nlm.nih.gov/bioproject/) under accession numbers OP169024–OP169026 (*A. sinensis*), OP169027–OP169032 (*C. yacare*), OP169033– OP169038 (*M. niger*), OP169039–OP169049 (*P. trigonatus*), and OP169050–OP169062 (*C. latirostris*).

## Acknowledgments

V.C.S.O was supported by the Coordenação de Aperfeiçoamento de Pessoal de Nível Superior, Brasil (CAPES) and Conselho Nacional de Desenvolvimento Científico e Tecnológico (CNPq) (401036/2022-7). M.A. was supported by the Charles University Research Centre programme 204069 and by Czech Science Foundation Project No. 20-27236J. M.d.B.C. was supported by the Conselho Nacional de Desenvolvimento Científico e Tecnológico (CNPq) (302928/2021-9). M.d.B.C. and T.L. were supported by Alexander von Humboldt Foundation (Research Group Linkage Programme). This study was financed in part by the Coordenação de Aperfeiçoamento de Pessoal de Nível Superior, Brasil (CAPES), Finance Code 001. We also would like to thank the Crocodile working group of the German Society for Herpetology and Terrarium Science association (DGHT AG Krokodile) for providing contact to keepers of alligators. We are grateful to Patrik Ferreira Viana, Hugmar Pains da Silva, Milena, Breno Almeida, Alexander Meurer, and Sebastian Scholz for all the valuable support provided with sampling.

## Institutional Review Board Statement

The study was conducted according to the guidelines of the Ethics Committee on Animal Experimentation of the Universidade Federal de São Carlos (Process number CEUA 4617090919). Collections were done under the authorization of the Chico Mendes Institute for Biodiversity Conservation (ICMBIO), System of Authorization and Information about Biodiversity (SISBIO-License No. 71857-7), and National System of Genetic Resource Management and Associated Traditional Knowledge (SISGEN-ABFF266). Blood samples from *Alligator sinensis* came from animals legally kept in Europe (CITES certificate number EU 0228-1057/14, ES-CC-00041/07C, ES-CC-00036/07C, 50721-18, DE-DA190814-5, DE-DA190814-6).

## Author contributions

M.B.C and R.U conceived the study. V.C.S.O., R.Z.S., C.A.G.G., R.M.C., M.A.G.R. and M.A., performed experiments. V.C.S.O., R.Z.S., C.A.G.G., R.M.C., M.A.G.R., M.A., T.E., T.L., F.P.., R.U and M.B.C analyzed the data. V.C.S.O., R.Z.S., R.U and M.B.C wrote the manuscript. All authors edited the final manuscript

## Supplementary Files

**Supplemental Table S1:** Similarity presented between the satellite DNAs of the same superfamily. Note that the percentage refers to the region in common between the two sequences.

| SatDNA | SatDNA | LOCAL Similarity (%) |
| --- | --- | --- |
| CyaSat01-41 | CyaSat04-41 | 70.78 |
| CyaSat01-41 | CyaSat06-286 | 79.0 |
| CyaSat03-60 | CyaSat05-200 | 73.4 |
| ClaSat01-41 | ClaSat06-1063 | 85.3 |
| ClaSat01-41 | ClaSat07-320 | 80.4 |
| ClaSat01-41 | ClaSat08-800 | 78.5 |
| ClaSat01-41 | ClaSat09-285 | 87.8 |
| ClaSat01-41 | ClaSat11-547 | 87.5 |
| ClaSat03-183 | ClaSat10-60 | 90.0 |
| ClaSat03-183 | ClaSat12-24 | 75.0 |
| MniSat01-41 | MniSat02-246 | 80.4 |
| MniSat01-41 | MniSat05-248 | 82.9 |
| MniSat01-41 | MniSat06-41 | 70.7 |
| PtrSat01-41 | PtrSat03-147 | 60.9 |
| PtrSat01-41 | PrtSat04-99 | 90.2 |
| PtrSat01-41 | PrtSat05-41 | 69.0 |
| PtrSat01-41 | PrtSat07-41 | 65.8 |
| PtrSat01-41 | PrtSat11-23 | 73.9 |
| PtrSat02-24 | PrtSat06-100 | 70.8 |
| PrtSat08-214 | PrtSat10-40 | 77.5 |
| ClaSat04-536 | PtrSat09-490 | 78.9 |

**Supplemental Table S2:** Summary of the BLAST searches against *Alligator sinensis* genome.

| Query id | Group | Number of alignments | Largest alignment (nt) | Number of distinct contigs |
| --- | --- | --- | --- | --- |
| CyaSat02-40 | 3 | 275 | 40 | 25 |
| ClaSat02-6317 |  | 6264 | 1073 | 500 |
| ClaSat04-536 | 4 | 150 | 443 | 49 |
| ClaSat05-40 | 3 | 481 | 40 | 41 |
| ClaSat13-398 |  | 1712 | 398 | 478 |
| MniSat04-40 | 3 | 124 | 40 | 15 |
| PtrSat08-214 | 3 | 28 | 210 | 9 |
| PtrSat09-490 | 4 | 133 | 387 | 45 |
| PtrSat10-40 | 3 | 214 | 40 | 18 |
| AsiSat01-1717 |  | 4602 | 1717 | 81 |
| AsiSat02-60 |  | 6700 | 60 | 43 |
| AsiSat03-96 | 3 | 287 | 96 | 23 |

**Supplemental Fig. S1.**
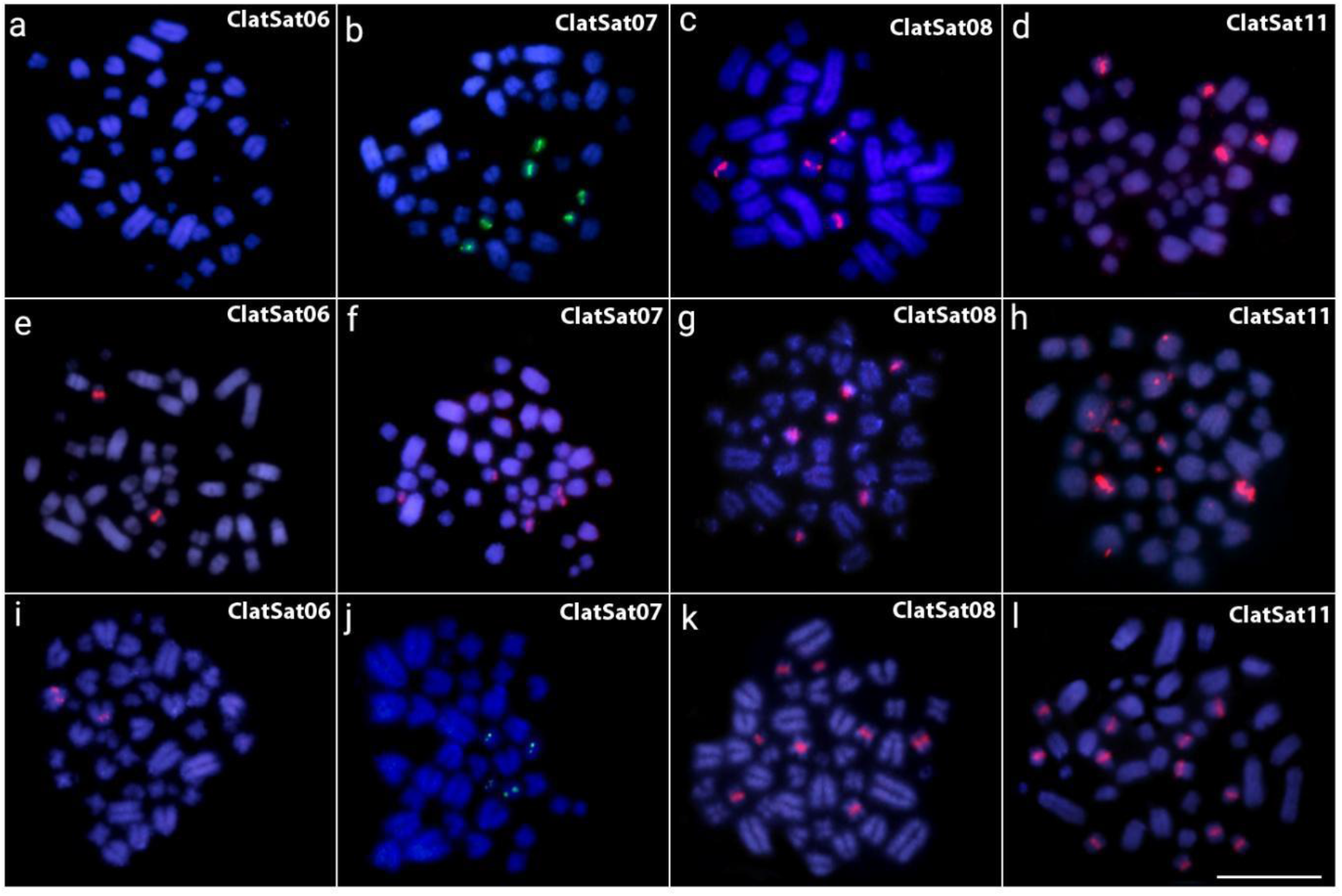
Metaphase chromosomes from *C. crocodilus* (a–d), *C. latirostris* (e–h) and *C. yacare* (i–l) after in situ mapping with satDNA probes belonging to group 1 (ClaSat06-1063; ClaSat07-320; ClaSat08-800 and ClaSat11-547). Chromosomes were counterstained with DAPI (blue). Scalebar=20μm

**Supplemental Fig. S2.**
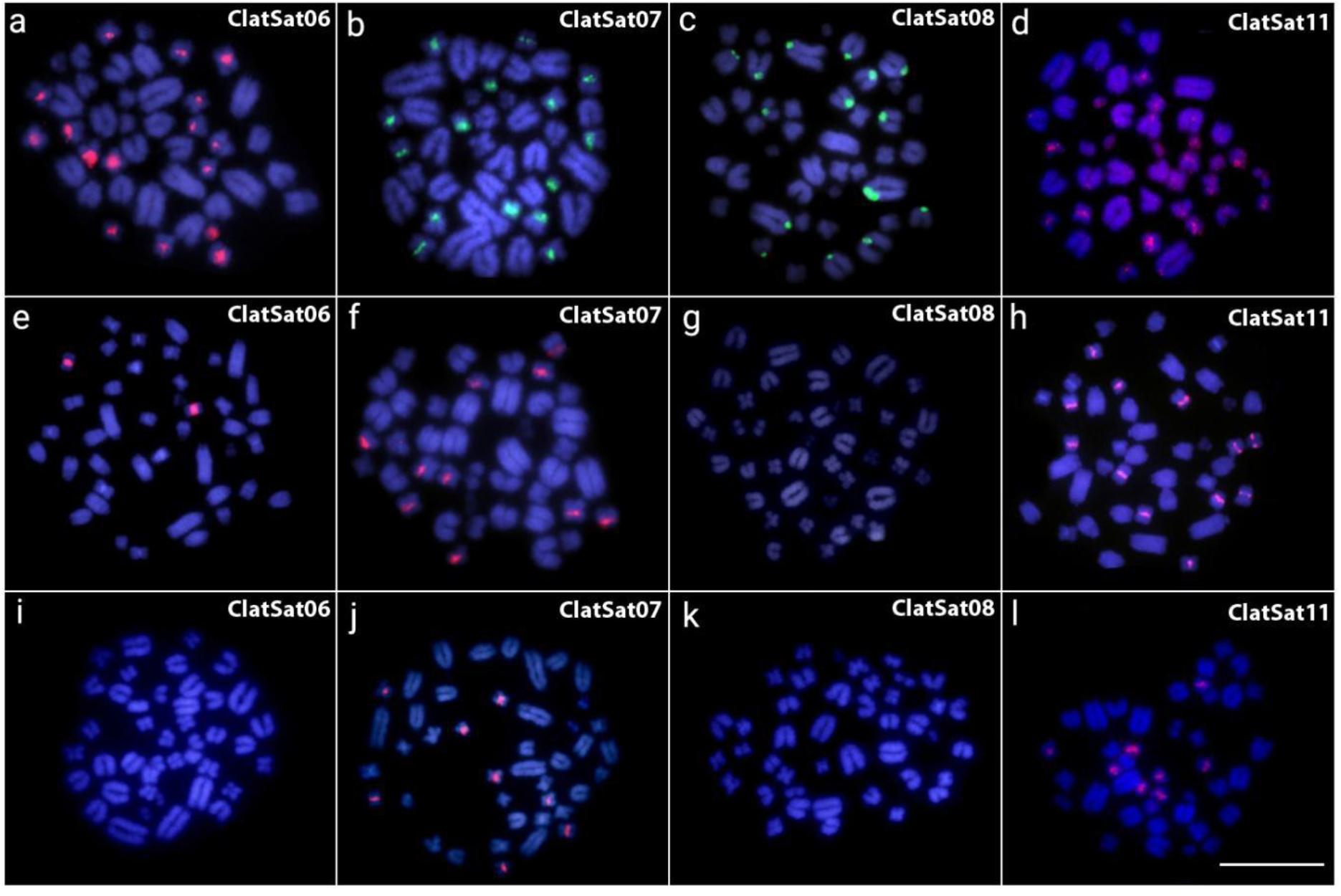
Metaphase chromosomes from *M. niger* (a–d), *P. palpebrosus* (e–h) and *P. trigonatus* (i–l) after in situ mapping with satDNA probes belonging to group 1 (ClaSat06-1063; ClaSat07-320; ClaSat08-800 and ClaSat11-547). Chromosomes were counterstained with DAPI (blue). Scalebar=20μm

